# Interconversion between Serum Amyloid A native and fibril conformations

**DOI:** 10.1101/2022.01.27.478066

**Authors:** Fatih Yasar, Miranda S. Sheridan, Ulrich H. E. Hansmann

**Affiliations:** Dept. of Chemistry & Biochemistry, University of Oklahoma, Norman, OK 73019, USA

## Abstract

Overexpression of Serum Amyloid A (SAA) can lead to a form of amyloidosis where the fibrils are made of SAA fragments, most often SAA_1−76_. Using Replica-Exchange-with-Tunneling, we study the conversion of a SAA_1−76_ chain between a the folded conformation and a fibril conformation. We find that the basins in the free energy landscape corresponding to the two motifs are separated by barriers of only about 2-3 *k*_*B*_*T*. Crucial for the assembly into the fibril structure is the salt bridge 26E-34K that provides a scaffold for forming the fibril conformation.

## 1 INTRODUCTION

Various illnesses, such as inflammatory bowel disease, rheumatoid arthritis, or certain cancers can cause overexpression of serum amyloid A (SAA) to concentrations 1000 times higher than what seen under healthy conditions. The resulting elevated risk for misfolding and aggregation leads in some cases to AA amyloidosis where damage from SAA-fibril deposits to tissues and organs adds to the symptoms, and complicates treatment, of the primary disease.^1–5^ For therapeutically intervention it is therefore necessary to understand the process of unfolding of the functional SAA structure and its re-organization into the conformation seen in the experimentally resolved fibrils.

Involved in cholesterol transport and the regulation of inflammation, the 104-residue long SAA forms hexamers.^6^ We have studied in previous work^7^ the conditions that lead to decay of the hexamer, cleavage of the released chains, and the subsequent misfolding of the resulting fragments (most commonly SAA_1–76_), which afterwards may associate into the SAA fibrils. We observed a competition between fast degradation of SAA fragments and the tendency of these fragments to form amyloids. Hence, in order to intervene, it is necessary to understand the kinetics of fibril formation. For this purpose, we study in the present paper now explicitly the conversion of SAA by modeling this process in all-atom simulations relying on a physical force field. As it is difficult for molecules of this size to obtain from regular molecular dynamics sufficient statistics, we utilize an enhanced sampling technique that was designed specifically by us for the investigation of structural tansitions. Our technique, Replica-Exchange-with-Tunneling (RET),^8–11^ allows us to observe the interconversion with sufficient detail to characterize important intermediates. Studying for a SAA_1−76_ fragment the transition between a conformation as seen in the folded and functional protein (PDB-ID: 4IP8),^12^ and the conformation seen in the experimentally resolved fibril structure (PDB-ID: 6MST)^13^ we find a characteristic sequence of events for this conversion that we relate to our earlier work and experimental measurements.

## 2 MATERIALS AND METHODS

### 2.1 Replica-Exchange-with-Tunneling

Deriving from computer simulations the mechanism by which SAA assumes its fibril form requires an exhaustive and accurate sampling of the free energy landscape of the protein. This, however, is a difficult task as typical time scales in protein simulations are only of order *µ*s. We have proposed in earlier work^8–11^ a variant of the Hamilton Replica Exchange method^14,15^ as a way to increase sampling for the special case of transitions between well-characterized states. For this purpose, we set up in our approach a ladder of replicas, each coupling a “physical” model with a structure-based model. On one side of the ladder the structure-based model drives the physical system toward the native state of SAA, while on the other side the bias is toward the fibril. The replica-depending strength of the coupling is controlled by a parameter λ that is maximal at the two ends, and zero for the central replica. Exchange moves between neighboring replicas lead to a random walk along the ladder by which the SAA conformation changes from one motif into the other. Accepting or rejecting these exchange moves with the criterium commonly used in Replica Exchange Sampling^16^ guarantees that the correct and unbiased distribution of the (unbiased) physical model will be sampled at λ = 0. However, the acceptance probability becomes vanishingly small for large systems. In order to avoid this problem we conditionally accept the exchange and rescale the velocities of atoms in the two conformations *A* (moving from replica 1 to replica 2) and *B* (moving from replica 2 to replica 1):

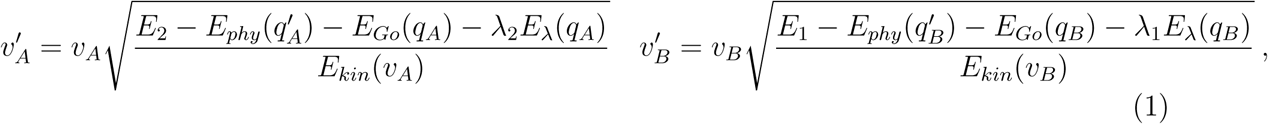

such that the total energy at both replica stays the same before and after the exchange. After the exchange, the systems on the two replica evolve by microcanonical molecular dynamics, exchanging potential and kinetic energy. Once the velocity distribution for each replica approaches the one expected at the given temperature, the final configuration 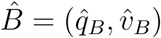 on replica 1 has a comparable potential energy to the configuration *A*, while the potential energy of 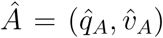 on replica 2 will be close to that of the configuration *B*. Defining 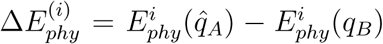, and accordingly 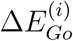 and 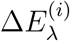, the *exchange is accepted with probability*

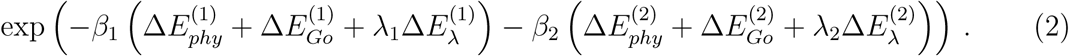

If rejected the simulation continues with configurations *A* (on replica 1) and *B* (on replica 2). We have shown that this *Replica-Exchange-with-Tunneling (RET)* procedure leads to the correct distribution for sufficient large systems and sufficiently long microcanonical segments, for detail see.^8,9^

### 2.2 Simulation Set-up

In order to study the interconversion of Serum Amyloid A (SAA) native and fibril conformations, we describe in our RET simulations the “physical” model of the system by an all-atom energy function, the CHARMM36m force field ^17^ in conjunction with TIP3P^18^ water molecules. The SAA_1−76_ fragment and 18,376 water molecules are put in a box of edge size 8.3 nm with periodic boundary conditions and the system neutralized with 2 sodium ions. The SAA fragment is capped with a acetyl group and a methylamine group on the N-terminus and the C-terminals, respectively. Start configurations are generated from the experimentally resolved structure of the full-size monomer as deposited in the Protein Data Bank (PDB-ID: 4IP8)^12^ by discarding residues 77-104, and randomizing the resulting fragment SAA_1−76_ in a 1 ns simulation at 1500 K, followed by another simulation of 1 ns at 310 K. After visual inspection for loss of secondary structure follows a final minimization of the resulting conformation. The biasing Go-model uses as target structures the fragment SAA_1−76_ as derived from the PDB-model 4IP8 after discarding residues 77 to 104, and the fibril model as deposited in the PDB under PDB-ID: 6MST.^13^ As the fibril model is only for the fragment SAA_2−56_ we had to add an arginine as first residues, and the C-terminal residues 57-76 by using CHIMERA.^19^ With the so-generated two target structures as input for the SMOG-Server^20^ at http://smog-server.org we derive then expressions for the corresponding Go-model energies E_*Go*_. Note that masses in the Go-models are scaled by a factor 14.21 to account for the lack of hydrogen atoms in the Go-models. Finally, we couple “physical” and Go-models by an energy E_λ_ as define in Ref.^21,22^ that quantifies the similarity between the two models. The strength of the coupling differs between the replicas. We used the following distribution of λ-parameters for the 24 replicas: λ = 0.08, 0.07, 0.06, 0.05, 0.04, 0.03, 0.02, 0.01, 0.007,0.006, 0.005, 0, 0, 0.005, 0.01, 0.02, 0.03, 0.04, 0.05, 0.06, 0.07, 0.08, 0.09, 0.1. Note that for the replicas 0-10 the coupling is between physical model and a Go-model is defined by the native conformation, while for replicas 13-23 the physical model is coupled with a Go-model defined by the fibril model. In order to avoid an odd number of replicas we use two central replicas with λ = 0 (replica 11 and 12), where the physical model is not biased by an Go-model.

The above set-up is implemented in a version of GROMACS 4.5.6,^23^ used by us to compare our data with earlier work. Hydrogen bonds are constrained for both the physical and go models using the LINCS algorithm^24^ leading to a time step of 2 fs. The van Der Waals cutoffs are set to 1.2 nm, and the Velocity Verlet algorithm^25^ is used to integrate the equations of motion. Temperature is controlled by the v-rescale thermostat.^26^ Note that our implementation of the RET algorithm requires different temperatures for each replica, which therefore slightly changes between 310 K and 310.23 K (in steps of 0.01 K). The length of the microcanonical segment is set to 1 ps in a RET move. Two independent trajectories of 100 ns and 85 ns were generated, taking measurements only from replica where λ = 0, i.e., where the physical model is not biased by any Go-term. Allowing for convergence of each trajectory we are left with a total of 125 ns for analyzing the free energy landscape of the system.

### 2.3 Analysis Tools and Protocols

We use GROMACS tools^23^ and the MDTraj^27^ software package in the analysis of the RET trajectories for measuring root-mean-square-deviation (RMSD), dihedral angles of residues, and number of contacts. Secondary structure analysis and evaluation of free energy landscapes is done by in-house-scripts. The transition pathway between native and fibril SAA conformations is derived from the free energy landscape projected on the RMSD to the respective structures. Using Dijkstra’s algorithm^28^ as implemented in the MEPSA software^29^ we construct for this purpose a minimum energy pathway between the two corresponding minima, allowing us to identify the barriers among different minima along the pathway. For visualization we use the PyMOL software.^30^

## 3 RESULTS AND DISCUSSION

We argue that the computational difficulties in simulating the conversion between native and fibril SAA conformations is alleviated in our variant of Replica Exchange Sampling. In order to support this assumption, we show in Figure 2a) the walk of a typical realization of our system along the ladder of replicas. At start time (*t* = 0) the physical system sits on a replica where it is biased with λ = 0.1 toward the folded structure, but walks numerous times between this replica and the opposite end-point of the ladder where the physical system is with λ_*max*_ = 0.08 maximally biased toward the fibril. The average exchange rate between neighboring replicas along the ladder is ∼ 25%. Monitoring the respective root-mean-square deviation (RMSD) toward both folded and fibril structure. we show in Figure 2b) that this walk through λ-space induces indeed interconversion between the two motifs.

**Figure 1:**
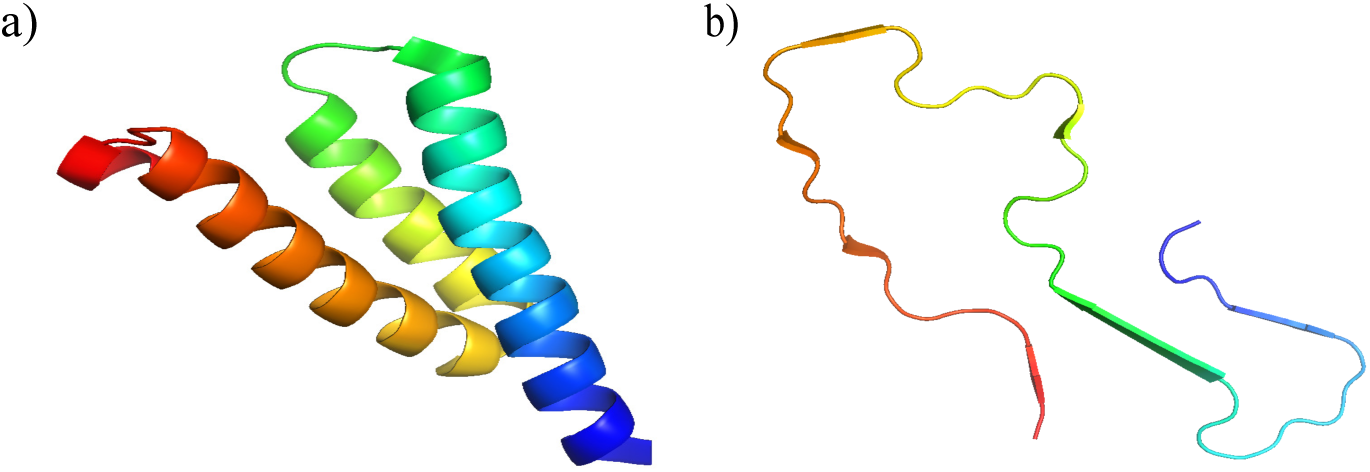
Structures of tSerum Amyloid A for the fragment SAA_1−76_ of the folded monomer (PDB-ID: 4IP8) (a), and for the chain in a fibril (segment SAA_2−55_, PDB-ID; 6MST) (b). The N-terminal residues are colored in blue.

**Figure 2:**
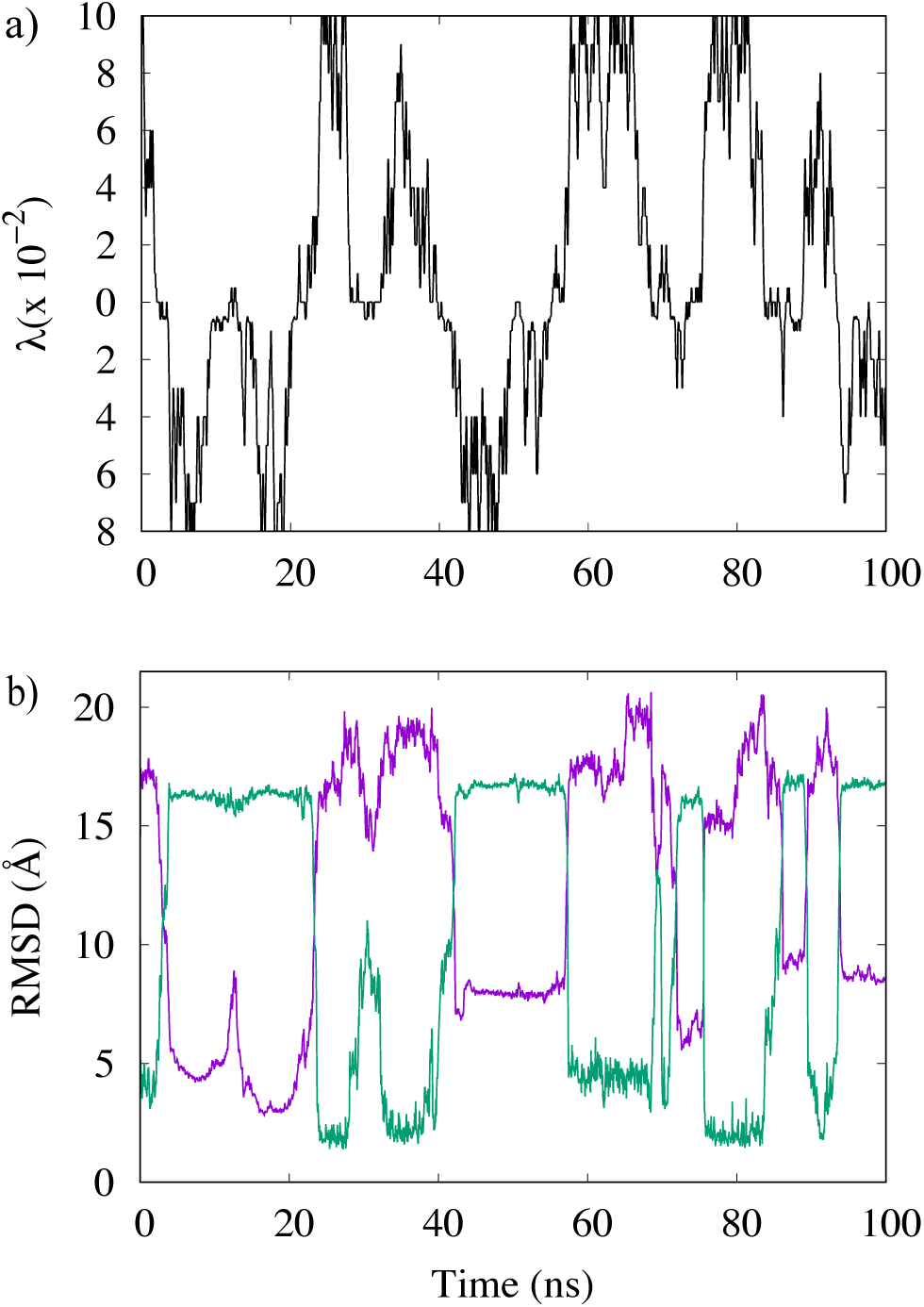
(a) A typical example of a replica walking through λ space starting from a replica, where the physical model is initially biased toward the folded SAA structure. While the system walks between replicas with bias toward the folded structure (upper half) and such with bias toward fibril structure (lower half), its configuration changes accordingly. This can be seen in b) where we show the corresponding time evolution of the RMSD to the native structure (in magenta) and the fibril structure (in green).

The higher rate of transitions allows for a more exhaustive sampling of the free energy landscape. A measure for the efficiency of our method, and a lower limit on the number of independent configurations sampled at the λ = 0 replica, is the number of walks across the whole ladder, from the replica with maximal bias toward the folded structure to the one with maximal bias toward the fibril, and back. The number of such tunneling events is inverse to to the average time needed to cross the ladder (termed by us the tunneling time). The higher the number of tunneling events, the shorter the tunneling time, and the more efficient will be our approach. This convergence of the simulation is checked by calculating comparing for two distinct time intervals. free energy landscape projected on the two distances introduced above. Resemblance of the data for these two intervals suggests that the simulation has converged after 10 ns, and therefore, we use the last 90 ns of this specific trajectory for our analysis. In this time span, we find at least four tunneling events with an average tunneling time of ∼ 27 ns.

While small, the existence of at least a few tunneling events gives us confidence in the reliability of our data at λ = 0, i.e., at a replica where the “physical” model of our system is not biased toward either structure. For instance, we show in Figure 3 the free energy landscape of the SAA_1−76_ chain projected on the root-mean-square-deviation (RMSD) to either folded or fibril conformation. This landscape is characterized by two prominent basins corresponding to either folded or fibril configurations. The interconversion process corresponds to a path connecting the two basins. We have shown in earlier work^31^ that the physical path does not necessarily corresponds to the ones seen in tunneling events as RET relies on an unphysical dynamic. Hence, in order to find a more realistic pathway, we have used the MEPSA software^29^ to determine the minimum energy pathway. This pathway, drawn as a black line in the landscape proceeds through a series of basins and appears to be thermodynamically reasonable. Note that by construction this minimum energy pathway does not connect specific configurations but bins. Each bin contains a certain number of configurations sampled throughout the simulations. These configurations may differ in their secondary structure or the native contacts seen in either the folded or the fibril structure. The frequency with which the three quantities are observed allows one to identify five distinct regions along the pathway that correlate with the basins and barriers of the landscape. These regions are labeled as A to E in Figure 3, and characteristic conformations for these regions are shown in Figure 4, where in the case of region A (characterized by a high frequency of folded conformations) and region E (rich in fibril-like conformations) these characteristic conformations are superimposed on the respective reference structures. Region C appears to be a transition region spanning a a large range of divers conformations. Note, however, that the barrier height is only between two and three *k*_*B*_*T*.

**Figure 3:**
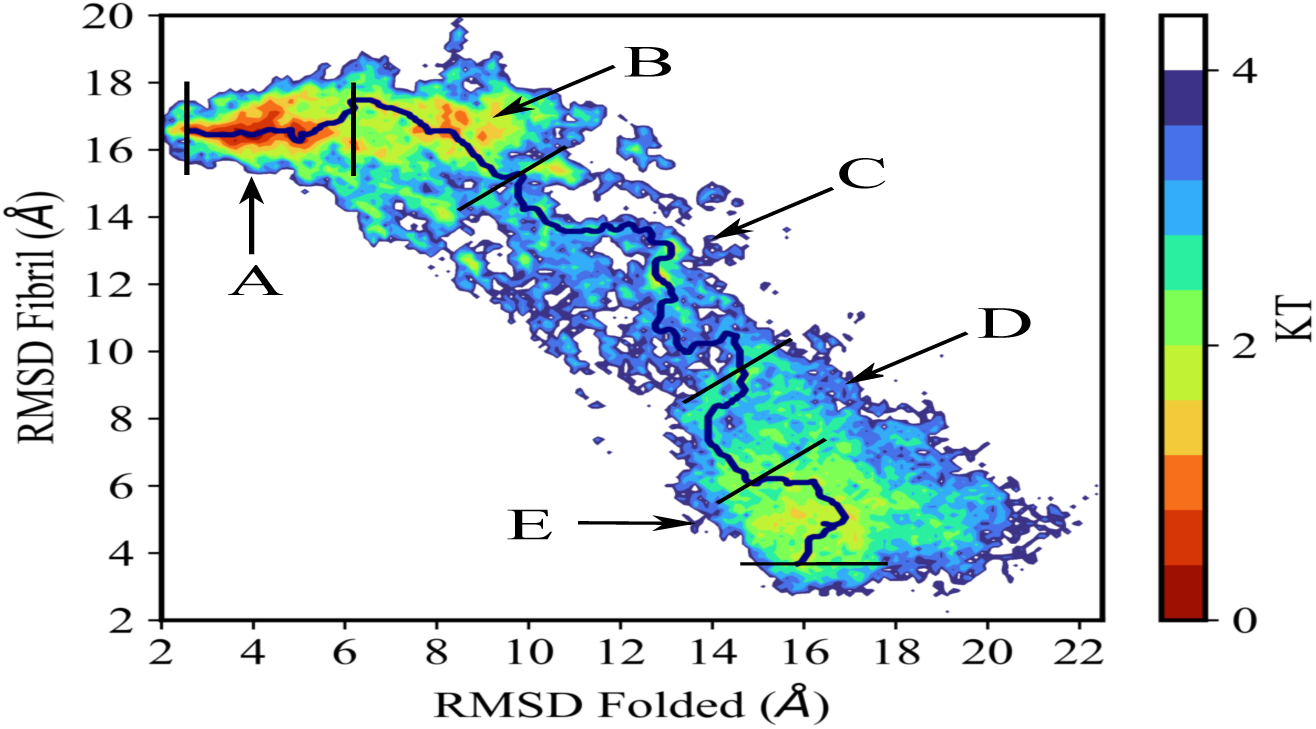
Free energy landscape as obtained from RET simulations, with data taken at λ = 0, i.e., where the physical models are not biased by any Go-term. Energies are listed in units of kT. The prospective transition pathway is drawn in black, and the five regions crossed by this path marked in capital letters.

**Figure 4:**
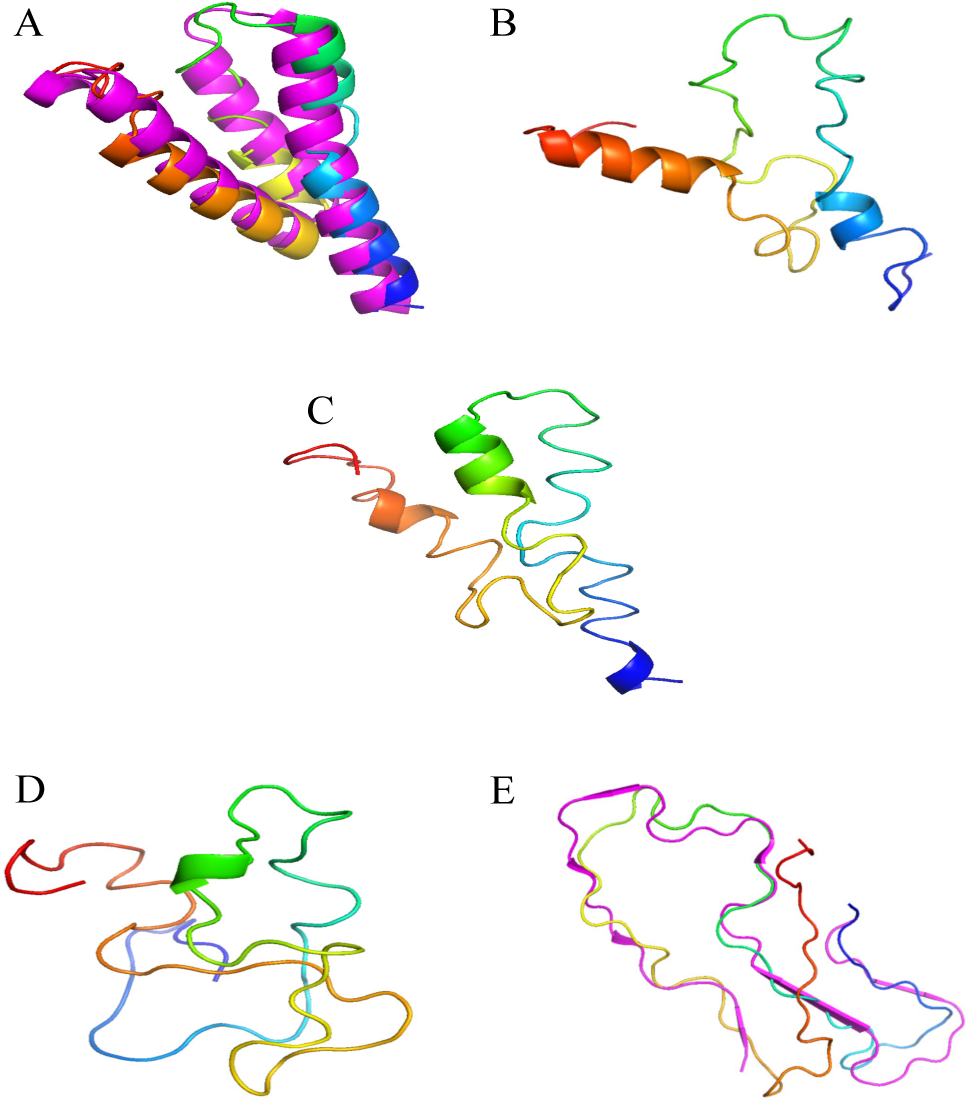
Characteristic conformations of of SAA_1–76_ as seen in each of the five regions identified on the proposed transition pathway. The N-terminus of the chains is colored in blue. For region A (dominated by folded-conformations) and region E (where fibril-like conformations are dominant) are these conformations superimposed on the respective reference structures.

In order to get insight into the structural changes along the proposed transition pathway, we augment the visual inspection of the dominant conformations by an analysis of the number *n*_*NS*_ of contacts found also in the folded structure, and of the number *n*_*FS*_ of contacts also found in the fibril structure. Here, we define a contact by the condition that the distance between at least one pair of heavy atoms in residues i and j is smaller than the cut-off value of the 4.5 Å. If in addition the two residues are separated by at least three other residues in the protein sequence, we call this a long-range contact. The two quantities, normalized to one, are shown in Figure 5a). As expected, the number *n*_*NS*_ of contacts found also in the folded structure decreases when going from region A, characterized by a low RMSD to the native structure, to region E (which has a low RMSD to the fibril structure). The opposite behavior seen for the number *n*_*FS*_ of fibril-like contacts. However, while there is a steep increase in *n*_*FS*_ when going from region C to region E, the corresponding change for the number *n*_*NS*_ of contacts seen also in the folded structure appears to be more gradually when going from region A to C. This picture changes when one considers only long-range contacts seen in the folded structure. This number *n*_*LR*_ is also shown in Figure 5a) and has a more pronounced behavior. While the number decreases rapidly going from region A to region B, and from region C to region E, it changes little between region B and C. This indicates that the conversion process from the helix-bundle of the folded conformations starts with decay of inter-helical contacts, while the transition region C is mainly characterized by the decay of intra-helical contacts, with the contacts found in strand-like conformations forming in region D. Note that we do not show separately number of long-range fibril-like contacts as by definition contacts involving strands are long-range.

**Figure 5:**
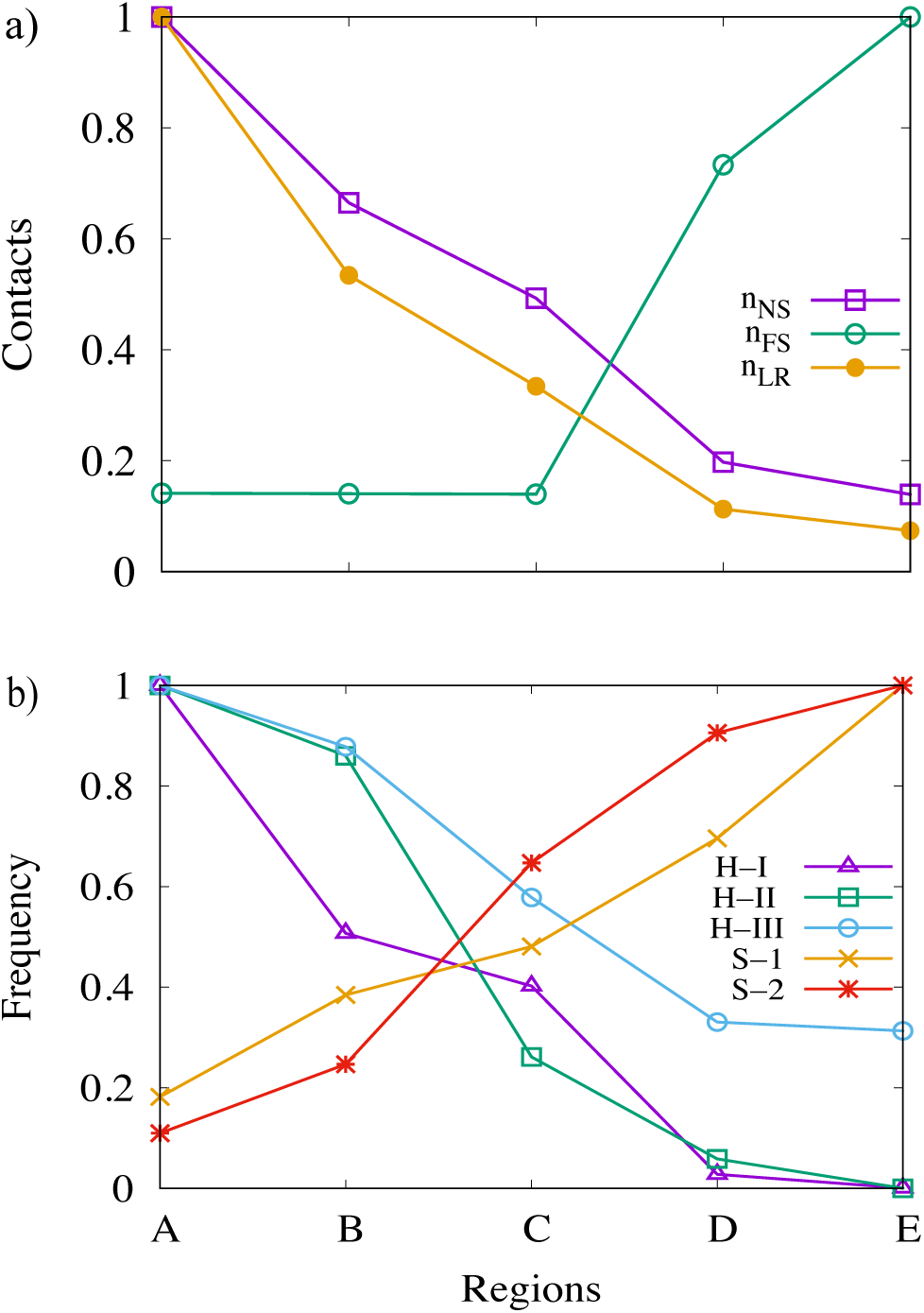
(a) The number of contacts (normalized to one) *n*_*NS*_ that are shared with the folded structure as measured in each of the five regions A to E of the transition pathway. The sub-set of long-range contacts *n*_*LR*_, again normalized to one, is drawn separately. Shown are also the number *n*_*FS*_ of contacts also found in the fibril reference structure. In (b) we show the relative frequency with which one of the three characteristic helices of the folded structure, or the two main *β*-strands of the fibril structure, are observed.

For a more detailed analysis we have also shown in Figure 5b the frequencies with which one of the three helices of the folded structure, or the two main strands of the fibril structure, are found in the five regions. Values are again normalized to one. The three regions of residues 2-27, 32-47, and 50-69 are considered helical if at least 30% of residues have dihedral angles as seen in an *α*-helix, that is if the dihedral angle pair (Φ, Ψ) takes values of between [-100^*o*^, -67 ^*o*^] and [-40^*o*^, -7^*o*^]. Similarly, we define the “strandness” of the two segments of residues 6-9 and 18-23 by the requirement that at least 20% of residues in the segment have dihedral angles as in a strand ([-160^*o*^, 160^*o*^] and ([100^*o*^, 100^*o*^]). The plots show that the N-terminal helix-I is the first to dissolve, with half of it already gone in region B. The central helix-II starts to decay later, but its propensity is also much decreased in the transition region C and neglectable in regions D and E. The C-terminal helix-III is the last one to decay and observed with substantial frequency even in region E. Note, however, that helix-III covers a region that was not resolved in the fibril structure and well may have transient helicity.

We observe a complementary picture for the two segment that are strand-like in the fibril. Again we have initially a larger frequency for the N-terminal segment of residues 6-9, with only in the transition region C the second segment gaining higher propensity for strand formation. Both the early decay of the first helix and the higher propensity of the N-terminal segment to form strands are consistent with experimental observations and our earlier work that demonstrated the importance of the N-terminal residues 1-11 and the role of helix-I for fibril formation.^7,32–35^ Interestingly, we observe in region B a mixture of the helix-broken and helix-weakened conformations discussed in Ref.,^7^ while in region C the conformation either lost their helicity or resemble more the aggregation prone helix-weakened conformations.

The above picture is also supported by the contact maps calculated for each of the five regions that are shown in Figure 6. Note, that the coloring does not indicate frequency of contacts but the average distance between residue pairs. As discussed in Ref.^7^ destabilization of the folded structure starts quickly after cleavage of the full-size monomer into a SAA_1−76_ fragment with a loss of contacts between residues 60-69 on one side, and residues 70-76 on the other side. Already in region A are only about 30% of these contacts still existing, and the frequency decreases further to about 17% in region B and C, before disappearing. As a result is helix-III more flexible than in the full-sized protein, allowing it to either break up (in helix-broken conformations) or partially dissolve (in helix-weakened conformations), thereby reducing contacts with helix-II. Overall is the loss of native contacts in regions A to C consistent with the process described in Ref.^7^ where the data were derived with a different simulation technique. On the other hand, in region D starts the formation of fibril-stabilizing intrachain salt bridges and contacts between residues 26 and 34, 26 and 46, 29-33, and 35 with 43 that has been also described in Ref.^34^ Note, however, that the contact between 29-34 in the helix-I - helix-II linker region is already observed with substantial probability in region A, and therefore likely crucial for fibril formation of SAA. Early formed is also the salt bridge between residue 26 and 34, seen in all five regions with about 10% - 20% probability. Hence, this contact is therefore a bottleneck/condensation point in fibril formation. Note that the contact 26-46 which competes with the 26-34 contact is seen in region D with 5% frequency but only with about 1% in the fibril region E. These results are also consistent with earlier work^36^ that emphasized the importance of the helix-I helix-II linker region for fibril formation, albeit we do not see the reported local unfolding of the two helices around this region.

**Figure 6:**
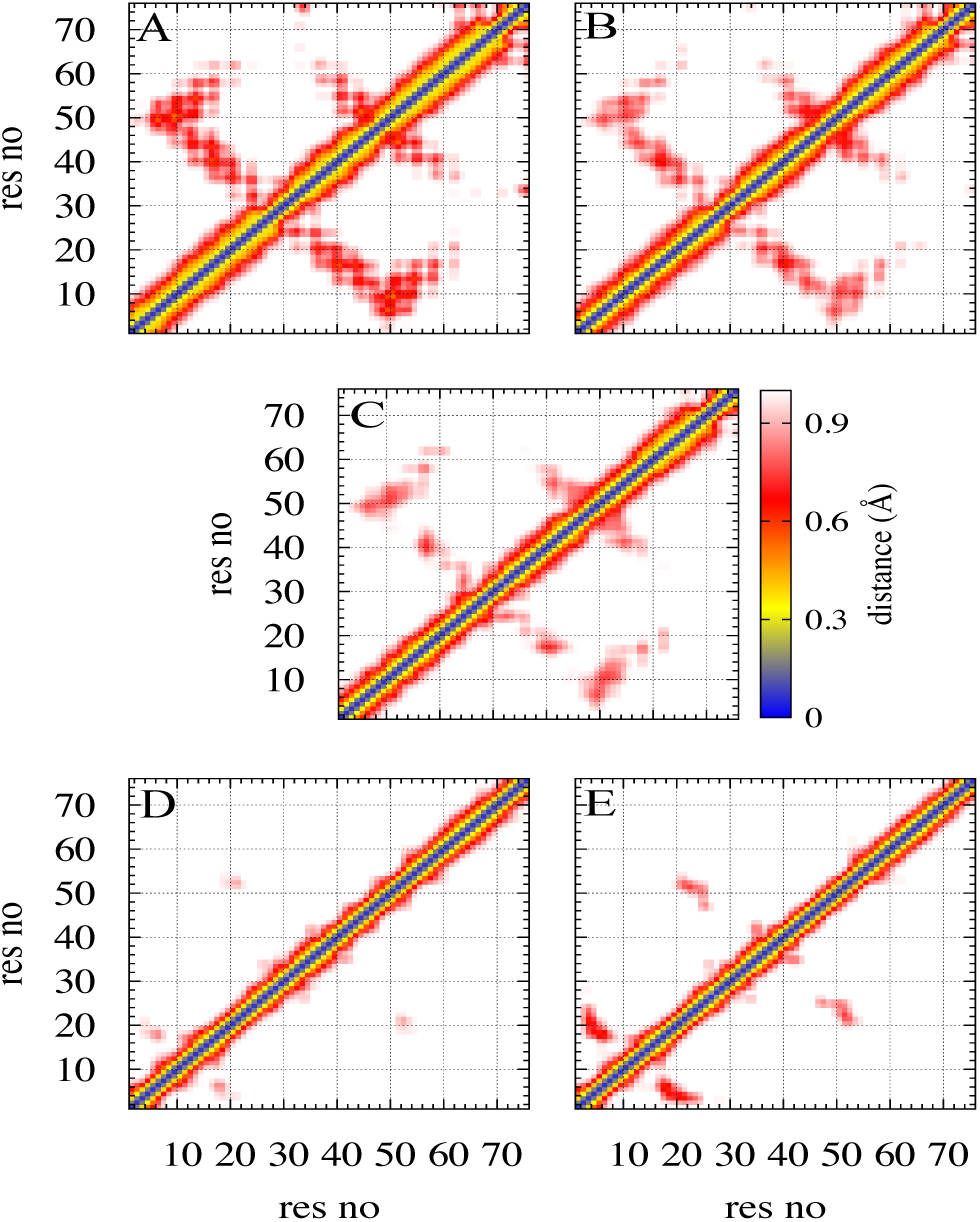
The residue-residue map of the average minimal distance between heavy atoms in a pair of residues, shown for each of the five regions. Residue pairs whose average contact distance is more than 10 Åare excluded. Numbers on the X and Y axis mark residues, and the average distance in Å is given by the color coding shown in the middle row.

## 4 CONCLUSIONS

Using a variant of Replica-Exchange-with-Tunneling (RET), we have probed the interconversion of SAA_1−76_ monomers between the folded structure and the one assumed in the experimentally resolved fibrils. *In vivo* this conversion happens at high (micro-molar) SAA concentrations, and likely will therefore depend on the interaction with neighboring chains. However, modeling the conversion under these conditions is not feasible, and we therefore restrict ourselves to an isolated monomer. Our assumption is that both folded and fibril configurations are prominent local minima in the free energy landscape of the protein, and connected by similar transition pathways as occurring under physiological conditions. The interaction with other chains becomes in this picture an external field that shifts the equilibrium towards the fibril structure. While this assumption is not proven, it appears to be reasonable and allows us to propose a conversion mechanism for SAA fibril formation, identifying critical residues and intrachain interactions, that may guide future experiments. Specifically, we find only small free energy barriers (of order 2-3*k*_*B*_*T*) separating folded and fibril structure that can be crossed easily once a critical nucleus for fibril formation is formed. Consistent with our earlier work ^7^ we find that the decay of the helix bundle of the folded structure progresses by a loss of inter-helical contacts between helix-II and helix-III and helix-I and helix-II, leading to the helix-broken or helix-weakened conformations of Ref.^7^Dominant on the pathway are the more aggregation prone helix-weakened conformations where the C-terminus of helix-III interacts with the C-terminus of helix-I and increases the flexibility of helix-I. The resulting higher entropy leads to de-stabilization of helix-I, causing a loss of helicity. Especially important is the release of the first eleven N-terminal residues from helix-I which then can misfold into strand-like configurations. This strand-segment is indeed the first one that appears in the conversion process seen in our RET simulation. We had shown in earlier work^34^ that the merging fibril structure is stabilized by intra-chain contacts between residues 26-34, 29-33 and 35-43, connecting residues located on the helix-I helix-II linker in the folded conformation. These contacts indeed appear once the barrier is crossed. Interestingly, the salt bridge between residues 26 and 34 is seen already transiently in early stages of the unfolding of the folded conformation and likely plays a crucial role for fibril formation. This could be tested by mutating one of these two residues to inhibit formation of this salt bridge.

## Acknowledgement

The simulations in this work were done using the SCHOONER cluster of the University of Oklahoma, XSEDE resources allocated under grant MCB160005 and TACC resources allocated under grant MCB20016 (both National Science Foundation). We acknowledge financial support from the National Institutes of Health under grant GM120634 and GM120578. FY also thanks to the Scientific and Technological Research Council of TURKEY (TUBITAK) under the BIDEB programs, and the Department of Chemistry and Biochemistry for kind hospitality during the his sabbatical stay at the University of Oklahoma.

## Notes

### Competing Interest Statement

The authors have declared no competing interest.

